# Sea star wasting disease in the keystone predator *Asterias rubens* from the Baltic Sea

**DOI:** 10.64898/2026.01.16.699924

**Authors:** Katja Seebass, Vinzent Ferfers, Jahangir Vajedsamiei, Frank Melzner

**Affiliations:** GEOMAR Helmholtz Centre for Ocean Research Kiel, Kiel, Germany; Christian-Albrechts-Universität zu Kiel, Kiel, Germany

**Keywords:** sea star wasting disease (SSWD)_1_, *Asterias rubens*_2_, Baltic Sea_3_, environmental stress_4_, marine disease ecology_5_

## Abstract

Sea star wasting disease (SSWD) is one of the most severe marine epidemics recorded, affecting numerous asteroid species and causing widespread population declines. Although a bacterial pathogen has recently been proposed for one species, its generality across taxa and regions remains unresolved. Here, we report the first year-round field assessment of SSWD in the Baltic Sea, a rapidly warming, low-salinity ecosystem hosting a single keystone sea star predator, *Asterias rubens*. Using field surveys, image-based monitoring, and laboratory experiments, we characterised disease dynamics and potential drivers in Kiel Fjord (western Baltic Sea). SSWD-like symptoms were present throughout 2024, with mean prevalence exceeding 40% across seasons and elevated levels during summer and early autumn, when sea surface temperatures approached 22 °C and salinity fluctuated between 11 and 17. The mean body radius of asymptomatic individuals declined from 5.6 cm in spring to 2.3 cm in early summer before partially recovering in autumn, consistent with high recruitment of juveniles and selective loss of larger, symptomatic individuals. In a complementary laboratory experiment, survival analyses identified body size as the strongest predictor of SSWD-associated mortality (hazard ratio = 50.8, *p* < 0.001), with large individuals far more likely to die than small ones. This size-selective mortality, together with environmental constraints on recruitment, suggests that SSWD may be reshaping population size structure and reducing predation pressure on blue mussels, with potential consequences for benthic community dynamics. Continued monitoring will be essential to assess the long-term impacts of SSWD on *A. rubens* populations and associated benthic ecosystems under ongoing climate change.

## 1 Introduction

The Sea Star Wasting Disease (SSWD) is one of the most devastating marine epidemics ever recorded, yet its cause remains largely unknown [1]. To date, more than 20 sea star species are affected by the disease that is characterized by skin lesions and ulcers, loss of body fluid pressure, loss of limbs and tissue degradation, often progressing rapidly to mortality [2–6]. In keystone species such as *Pisaster ochraceus* and *Pycnopodia helianthoides*, both highly affected by the disease, populations declined by up to 90%, triggering regime shifts from macrophyte-dominated to mussel- and barnacle-dominated ecosystems [7–9]. Initially confined to the Pacific, the disease spread to the Atlantic in 2016 [10] and was first documented in European waters in 2022, affecting captive *Crossaster papposus* and *Asterias rubens* collected from the Irish Sea [11,12] as well as captive *A. rubens* from Helgoland [13]. This progression suggests that limited field monitoring in Europe may have delayed detection in wild populations, underscoring the need for increased surveillance to predict a further spread of the disease.

Further, despite more than a decade of research, the exact cause and transmission pathway of SSWD remain unclear [1]. Early studies suspected a viral agent (‘Sea Star associated Densovirus’, SSaDV) [14], but this hypothesis was later ruled out, as the virus appears to be a natural component of the microbiome in several sea star species [15,16]. Subsequent work has documented progressive microbial dysbiosis with symptom onset, often involving increased abundance of potentially pathogenic and copiotrophic bacteria, such as *Vibrio* spp. [17–21]. Most recently, the hypothesis of a specific bacterial pathogen has gained renewed support through the isolation and identification of *Vibrio pectenicida* strain FHCF-3 in diseased *P. helianthoides*, which fulfilled Koch’s postulates in experimental infections [22]. This strain, closely related to a known *Pecten maximus* pathogen (Lambert et al., 1998), which produces a haemocyte killer toxin that suppresses host immune responses, was especially enriched in the coelomic fluid of diseased *P*.*helianthoides* [22]. These findings represent the strongest mechanistic evidence to date for a bacterial etiology of SSWD and suggest that infection may proceed via invasion of the coelomic cavity [22,23].

However, given the consistent observation of microbial dysbiosis and the involvement of multiple potentially pathogenic taxa, it remains possible that SSWD stems from a polymicrobial infection rather than a single-pathogen disease [1,19,21]. Host susceptibility may also be modulated by physiological condition and environmental context - for example, immune suppression following spawning which has been linked to increased disease vulnerability in other marine invertebrates, such as oysters [24–26]. Temperature has also emerged as a potential driver of SSWD dynamics, with elevated and variable thermal conditions frequently coinciding with disease outbreaks in field observations [7,8,27–29]. Since temperature influences a range of ecological and physiological parameters, including oxygen availability and primary productivity, it may act both directly and indirectly on disease processes [17,30]. In contrast, some cooler environments appear to buffer against widespread mortality, possibly acting as thermal refuges for vulnerable species [31]. Importantly, current evidence for *V. pectenicida* FHCF-3 as a causative agent remains geographically and taxonomically restricted to Pacific *P. helianthoides*. Whether this bacterium plays a comparable role in other sea star species - including those already affected in the North Atlantic - remains to be seen [32]. Nonetheless, given that SSWD likely spreads via ocean currents, and that *Vibrio* spp. species are highly responsive to environmental conditions, the potential for continued expansion into vulnerable marine ecosystems is substantial [33]. As disease surveillance in marine systems remains limited, outbreaks may go undetected until mass mortality events occur [34,35].

Since its discovery in the North Sea by Smith and colleagues in 2022, one of the main concerns has been the potential spread of SSWD from the North Sea into the Baltic Sea via the Danish Straits (Smith et al., 2022). The Baltic Sea is an ecologically vulnerable region undergoing rapid warming, with temperatures rising at +0.6°C per decade since 1959 and projected to increase by +2°C by the end of the century [36,37]. Additional stressors, including eutrophication, low salinity, and expanding hypoxic zones, further threaten ecosystem stability and functionality [38–41]. Compared to more biodiverse marine systems, the Baltic Sea has relatively low species diversity, making it more susceptible to cascading ecological effects if key species are lost [42–44]. One such key species is the sea star *Asterias rubens*, the only predominant asteroid echinoderm in the western Baltic Sea. As a benthic predator, *A. rubens* plays a crucial role in maintaining ecosystem stability by regulating populations of blue mussels (*Mytilus edulis × trossulus*) [45]. Without this predation pressure, mussel populations may expand unchecked, outcompeting other important ecosystem engineers such as eelgrass (*Zostera marina*), which is essential for sediment stabilization and habitat provision [46,47].Experimental studies indicate that *A. rubens* is already highly vulnerable to Baltic-specific stressors, including salinities below 12 and elevated temperatures during both summer (>22°C) and winter [48–52]. These continuous environmental pressures highlight the particular risk that SSWD poses to Baltic *A. rubens* populations. Yet, no systematic monitoring of wild populations has been conducted in this region so far.

To address this gap of knowledge, we carried out a systematic, camera-based survey of western Baltic *A. rubens* populations in Kiel Fjord, assessing sea star abundance, signs of disease, SSWD symptom prevalence, and mortality. We document, for the first time, the occurrence of SSWD-like symptoms (hereafter SSWD) in the Baltic Sea, thereby establishing essential baseline data for tracking its progression. By following the population over the course of one year, this study provides insights into the temporal development and manifestation of SSWD in the field. In addition, we investigated potential physiological factors influencing disease dynamics, focusing on body size. For the laboratory experiment, sixty sea stars were collected from Kiel Fjord, classified into a size class, and monitored over 33 days to test whether body size affects the progression and severity of SSWD. Complementary to this, up to 40 individuals were collected monthly from the field, classified, measured, and sampled for coelomocyte and tissue analyses. Together, these efforts provide the first assessment of SSWD symptoms in Baltic *A. rubens*.

## 2 Methods

### 2.1 Monitoring procedure

Weekly field monitoring began in January 2024 to assess *A. rubens* abundance and SSWD symptoms within naturally occurring blue mussel (*Mytilus* spp.) reefs in Kiel Fjord (Western Baltic; 54°19′45″N, 10°08′56″E). Surveys were conducted at four fixed stations (“Stations” 1-4), each consisting of a central bridge and two floating docks (left and right transects, ~21 m each, depth 1.1–3.7 m). The reefs at these stations ranged from 735 to 1080 m^2^ in area (mean ≈ 900 m^2^) and approximately 80-100 m^2^ were recorded per reef per survey (~10 % of reef area; ~360 m^2^ total per week). Video footage was collected using GoPro Hero 10 cameras (GoPro Inc., California) mounted on a 3.9 m adjustable pole and positioned 95.5 cm above the substrate, capturing ~1 m^2^ per frame (calibrated with 50 cm scale bars, later removed). Along each transect, sequences consisted of two frames (Frame 1 at the jetty edge, Frame 2 one-meter outward; Supplement), followed by repositioning 1.2 m along the transect. This procedure yielded ~20 short video clips (~40 frames) per transect, covering ~40 m^2^ per side. Videos were recorded in 4K/30 fps (linear mode) to allow detailed identification of SSWD symptoms. At each session, temperature and salinity were recorded.

#### 2.1.1 SSWD classification framework

To characterize disease progression in *A. rubens* from the Baltic Sea, we developed a modified classification framework for sea star wasting disease (SSWD). The commonly used four-to five-stage model, typically describing lesion initiation on a single arm or the central disc, followed by progressive spread [7,14,18,20,21] did not capture the frequent multilocal and variable onset observed in our field and laboratory observations. Our criteria were formed by previous studies and our own observations in *A. rubens* [12,13].

Early signs such as arm curling (“ray dysplasia”; [1,22]) and swelling [5] were often accompanied by localized hydrostatic bulging [8,53], papulae retraction, and oedema, possibly with localized lesions occuring [5,13]. These presentations were grouped into a mild category (approximately equivalent to Stage 1, Figure 1c & c.i). An advanced category (Stage 2-3 equivalent) included individuals showing generalized discoloration, pale surface lesions, softening of the integument, and reduced attachment strength (Figure 1d-d.i). Mortality can occur in this stage, prior to arm autotomy. The moribund category (corresponding to Stage 4 in some definitions) included individuals with everted stomachs or open ulcers, particularly at arm-disc junctions, together with major loss of turgor and body fluids, although tube-foot movement could still be present but reduced (Figure 1e-e.i). Individuals exhibiting black necrotic tissue, immobility, retracted tube feet, and inverted posture were classified as dead. Rapid, simultaneous arm loss was rare but noted separately. Individuals with black necrotic tissue, immobility, retracted tube feet, and inverted posture were considered dead.

**Figure 1.**
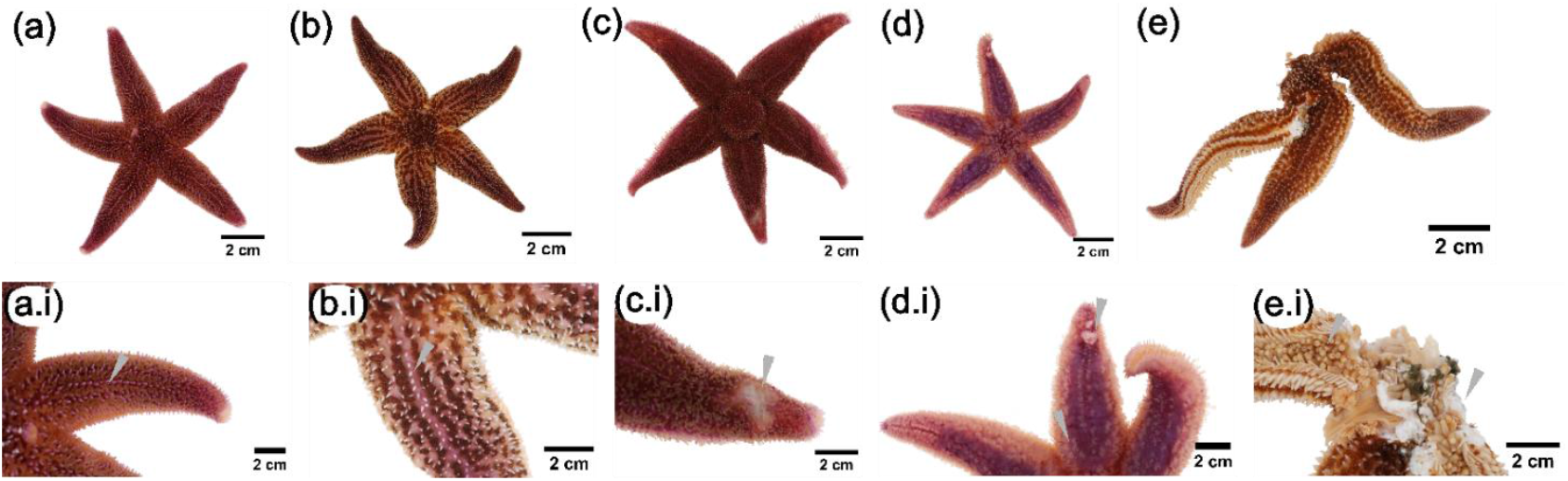
Examples of phenotypes and symptoms of sea star wasting disease in *A. rubens* from the Baltic Sea, ranging from (a, b) asymptomatic to (e) moribund or dying individuals. Lower panels (a.i-e.i) show close-ups of the corresponding sea stars. (a) Smaller asymptomatic individual with homogeneous integument, intact papulae, and uniform coloration. (b) Asymptomatic individual of larger body size. The ossicles may spread, particularly when the gonads have ripened, producing an oedema-like appearance that is normal for *A. rubens* in the Baltic Sea. (c) Individual showing mild symptoms, including skin softening, slight arm curling, and a localised pale lesion (c.i, arrow). (d) Advanced stage with retracted papulae, discolorations covering >50 % of the epidermis, pale patches and open lesions (d.i, arrows); mortality may occur at this stage before arm autotomy. (e) Moribund individual exhibiting severe tissue and fluid loss, partial body-wall disintegration, and only limited tube-foot movement. Extensive tissue and fluid loss lead to death within 24-48 h in captivity.

#### 2.1.2 Downstream analysis in BIIGLE

Two frames per short video (Frame 1, Frame 2) from left transects (~21 m) were extracted and uploaded to BIIGLE (biigle.de; [54]) under the project Kiel_Fjord_Monitoring. Stations 1-2 and 3-4 were filmed on separate days each week. To capture seasonal variation, a subset of seven weeks in 2024 was annotated (Winter: 14/19 Feb, 11/15 Dec; Spring: 12/15 Apr; Summer: 12/15 Jun, 11/16 Jul; Autumn: 16/19 Sep, 11/15 Oct). The annotation library included Asymptomatic, Mild, Severe (now referred to as Intermediate), Moribund/Dead, and SeaStar_No_ID (for individuals only partially visible or blurred). Additionally, three random frames per date and station were selected to measure sea star radius of all individuals in the frame. A 50 cm scale bar served as reference. Measurements were performed in BIIGLE using line strings (pixels) and converted to centimetres with a fixed scale factor (50 cm = 1346.5 px). Dot annotations and line-string data were exported as CSV files for analysis in R. Records with implausible sizes (< 1 cm or > 15 cm) were excluded. For descriptive visualization, individuals were grouped by health status (asymptomatic, mild, or other), and body-radius distributions were plotted to illustrate seasonal patterns. No formal statistical tests were applied at this stage.

#### 2.1.3 Animal collection and sampling method

Each month, up to 40 *A. rubens* were collected by snorkelling around the filming transects, alternating between stations to avoid oversampling. Except for January, June, November, and December (see Table S1), 20 individuals (10 asymptomatic, 10 symptomatic) were collected biweekly and transferred individually (1.2 L aquaria) to GEOMAR, Kiel. Upon arrival, individuals were photographed, and arm length (center to arm tip) and wet weight were recorded. Sea stars were staged according to disease symptoms, then sampled for coelomic fluid and tissues. Approximately 2 ml of coelomic fluid were withdrawn from one arm tip using sterile scissors and flash-frozen in liquid nitrogen. Tissue samples (~0.5 cm) were excised from three arms (one lesion and two unaffected sites, or three unaffected sites) along with pyloric caeca and, when possible, gonad tissue, and all were flash-frozen for downstream analyses.

### 2.2 Laboratory survey

#### 2.2.1 Animal collection and experimental setup

On 22 May 2025, 60 *A. rubens* were collected from Stations 2 (54.32889° N, 10.14861° E) and 3 (54.32917° N, 10.14889° E) in Kiel Fjord. Surface water temperature was 16 °C and salinity 13.6. To prevent cross-contamination, individuals were collected separately using 1 L plastic bags, which also served as gloves. Sea stars were categorized by health status and size: small (<4 cm), medium (4–6.5 cm), and large (>6.5 cm). Health categories included Asymptomatic (no visible signs of SSWD) and Diseased (SSWD-like symptoms such as lesions, arm twisting, bloating, or arm loss). Due to the high prevalence of SSWD, healthy individuals >6.5 cm were unavailable, resulting in unbalanced health categories and a strong size-disease association. Wet weight was recorded within collection bags to avoid contamination. Upon arrival at GEOMAR climatized chamber (17 °C), individuals were re-examined, scored for disease status (Table X, supplement), and placed in individual 12.5 L Plexiglas aquaria (10 L ASW, salinity = 15). The experiment was originally designed with three treatment groups: Control, Diseased, and Exposed. Diseased individuals were naturally symptomatic at collection, while Exposed individuals (n = 11) were initially asymptomatic but received a mixture of seawater from tanks that had contained recently deceased, symptomatic sea stars. However, this exposure treatment had no measurable effect (only one Exposed individual developed symptoms) and all large and many medium-sized individuals from the Diseased group died rapidly despite separate housing. As a result, subsequent analyses focused on identifying predictors of survival across individuals rather than on treatment effects. Within 24 h, 13 of 60 sea stars spawned, likely due to handling stress or environmental change; water in all aquaria was subsequently replaced. This spawning event was considered in analyses of disease progression and survival. Originally, the study aimed to test infection dynamics using 40 healthy and 20 diseased individuals, but high field prevalence and rapid mortality among large sea stars led us to focus on natural disease dynamics and size-dependent susceptibility. All aquaria were continuously aerated using air stones, and 2 L of water were replaced daily with freshly prepared artificial seawater (ASW; salinity 15.0, 17 °C). Gloves were sanitized with 80% ethanol or changed between animals, and separate equipment was used for diseased and asymptomatic individuals to prevent cross-contamination. Sea stars were inspected daily for posture, body surface, and tissue integrity, and SSWD stage was updated accordingly. Disease progression was documented with frequent photographs taken using an underwater camera. Individuals with arm autotomy or found dead were removed, photographed, and dissected. Tissue samples were flash-frozen in liquid nitrogen and stored at −70 °C. After day 33, all remaining individuals were dissected and preserved.

### 2.3 Statistical analysis

#### 2.3.1 Field data

All statistical analyses were conducted in RStudio (v4.3.3; R Core Team, 2025). Data processing and visualization used the tidyverse [55], lubridate [56], and ggplot2 packages [57]. To visualize seasonal variation in disease prevalence, we fitted preliminary generalized linear mixed models (GLMMs) to aggregated site-level data. These models were used to visualise temporal trends rather than for formal statistical inference. For each sampling date and station, the proportion of affected individuals (either showing mild signs of SSWD, or classified as moribund = severe + mortally ill) was modelled using a beta-binomial error distribution with a logit link. Time was included as a smooth term (bs(t_center, df = 4)), and station identity was included as a random intercept to account for repeated sampling. Models were fitted using the glmmTMB package [58]. Sea star density was modelled using an analogous negative-binomial GLMM with a temporal spline and station as a random effect. Model fit was assessed using simulated residuals from DHARMa, and predictions with 95 % confidence intervals were extracted for plotting. Model diagnostics indicated no problems with residual distribution, dispersion, zero-inflation, or overall model fit.

#### 2.3.2. Laboratory survey

Differences in body size (arm length cm, weight g) across treatment groups were tested using one-way ANOVAs with Tukey’s HSD post hoc tests. Survival differences among groups were analysed using Kaplan-Meier curves and log-rank tests using the survival package; [59]. To evaluate effects of treatment, body size (length, weight), and spawning status on mortality risk, Cox proportional hazards models were fitted and visualised using the survminer package [60].

## 3 Results

### 3.1 SSWD symptom progression shows some variability

Disease progression was not strictly linear. Some individuals died during the advanced stage without displaying all features typical of moribund animals, such as extensive body-wall disintegration. Arm autotomy was inconsistent: while it occurred in a subset of individuals, several clearly moribund or dead sea stars retained all arms. Occasionally rapid, near-simultaneous loss of multiple arms was noted (1-2 out of 60). Dead individuals were typically characterized by black necrotic tissue, complete immobility, retracted tube feet and an inverted posture, indicating that mortality can occur at multiple points along the progression rather than exclusively at the most extreme stage.

### 3.2 Dynamics of disease prevalence throughout the year 2024

Analyses were based on a subset of 1,131 annotated frames (≈1 m^2^ each) collected from four fixed stations across seven seasonal time points in 2024. In total, 19 860 *Asterias rubens* were recorded; because transects were revisited over time, this number reflects cumulative sampling effort rather than unique spatial coverage. Disease prevalence showed clear seasonal variation, with predicted SSWD levels ranging between ~40–55% (45.9% on average) and moribund prevalence between ~10-25% (Figure 2). Over all time points analysed, 3250 individuals were observed in terminal disease categories (severe or moribund) within the annotated transect subset, which represented ~5 % of the total reef habitat (videos covered ~9 %, but only half of all frames were analysed). Environmental conditions varied strongly over the year. Seawater temperature followed a unimodal seasonal cycle, peaking in August, whereas salinity decreased sharply in spring and early summer before rising again in autumn (Figure 2). Peak summer temperatures approached the 22 °C stress threshold for *A. rubens*, and salinity occasionally dropped near 11 during spring and autumn, coinciding with frequent upwelling events in the Baltic Sea.

**Figure 2.**
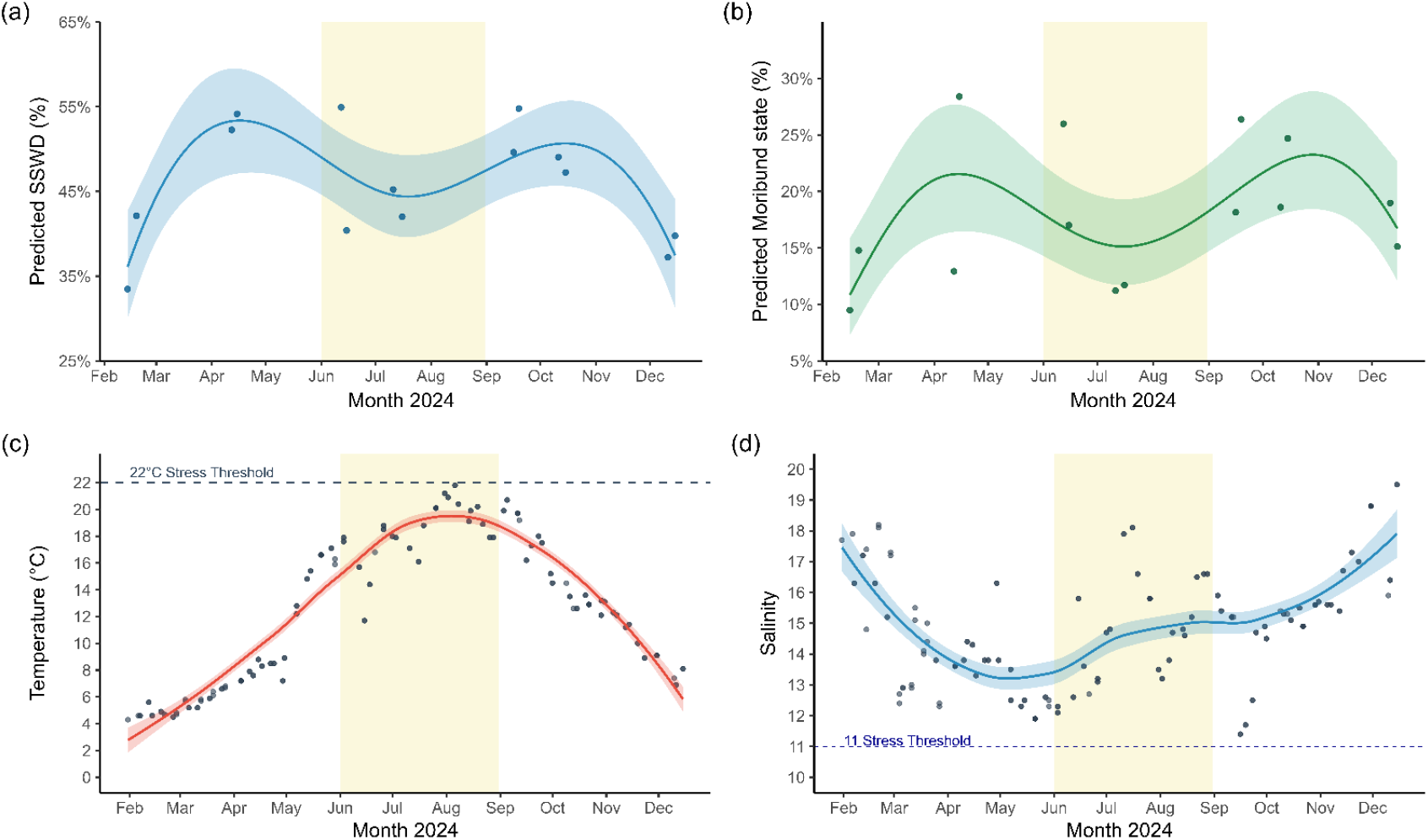
Seasonal patterns of sea star wasting disease (SSWD) in *A. rubens* and environmental conditions in Kiel Fjord in 2024. (a) Predicted prevalence (%) of individuals showing visible SSWD symptoms, based on video monitoring of field populations. A beta–binomial mixed model revealed a bimodal seasonal pattern with peaks in spring and autumn (*p* < 0.01). (b) Predicted proportion (%) of moribund individuals (near death or dead) showing a similar pattern (*p* < 0.01). (c) Sea surface temperature (1 m depth) with the 22 °C thermal stress threshold indicated by a dashed line; peak summer temperatures approached but did not exceed this threshold (maximum = 21.7 °C). (d) Seasonal variation in salinity, with a dashed line marking the stress threshold of 11 for Baltic *A*.*rubens*. The summer period (June-August) is shaded in yellow for reference. Shaded ribbons in all panels represent 95 % confidence intervals.

### 3.3 Seasonal variation in body size, population density and mortality dynamics in 2024

Across all sampling dates, mean body radius of *A. rubens* ranged from 1 cm to 14.7 cm, with an overall mean of 3.9 ± 2.2 cm (mean ± SD). When grouped by disease status, asymptomatic individuals averaged 3.9 ± 2.2 cm (n = 841), mildly affected individuals 3.8 ± 2.2 cm (n = 675), and moribund individuals 3.4 ± 1.8 cm (n = 399). Asymptomatic individuals generally followed the same seasonal pattern in size variation as the total population (Figure 3 b). Across all stations, mean body size decreased during early summer while estimated sea star density increased over the same period, followed by a decline towards autumn. Sea star density varied significantly over time (χ^2^= 43.17, df = 4, p < 0.001), with higher densities in early summer and lower densities toward autumn (Figure 3 c). Mean density was 17.0 ± 18.4 individuals m^−2^.

**Figure 3.**
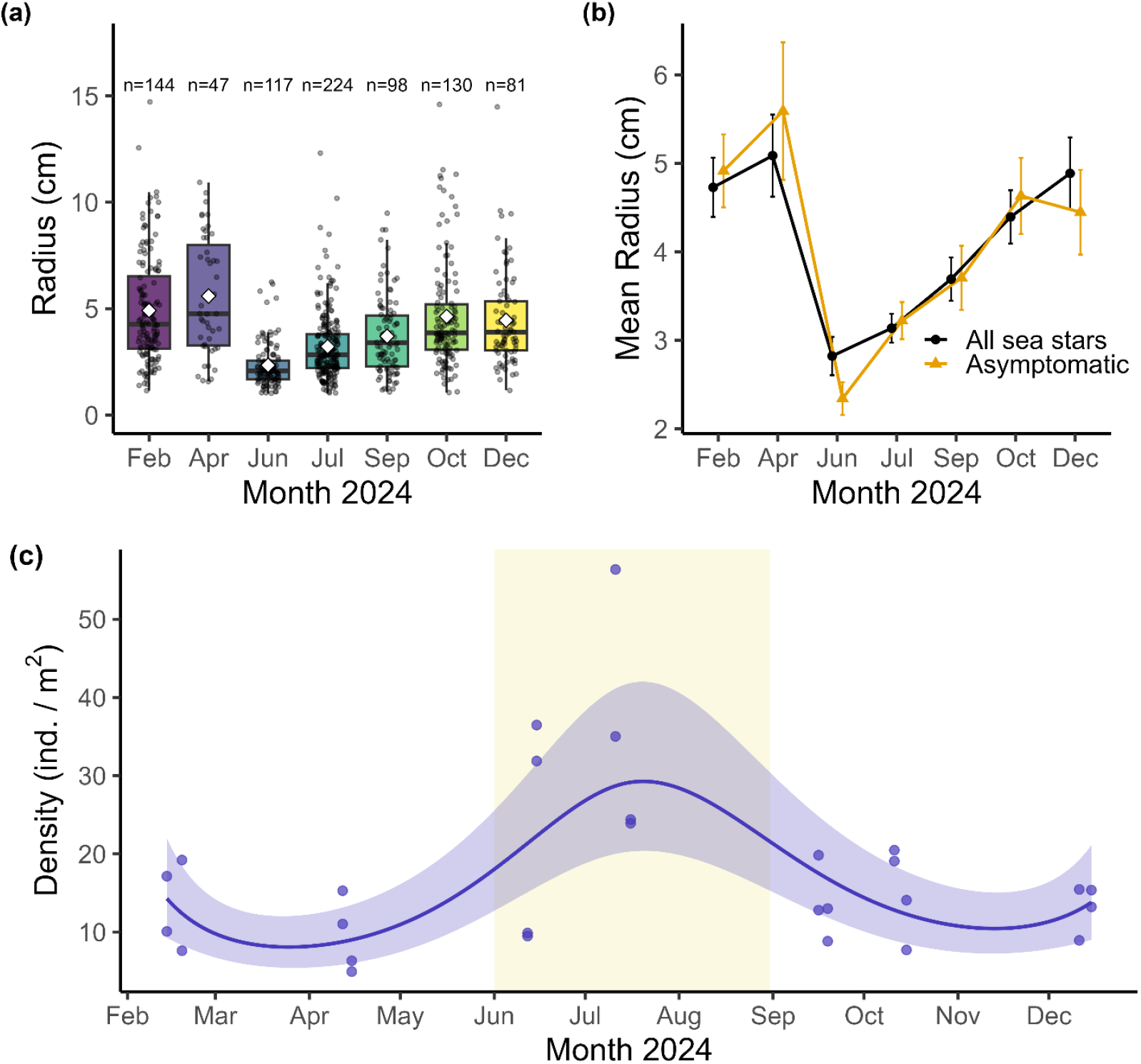
Seasonal variation in body size of *A. rubens* in Kiel Fjord in 2024. (a) Radius (cm) of asymptomatic sea stars measured from three annotated frames per date and station in BIIGLE, using 50 cm scale bars for reference. Median radius was smallest in summer (June-July) and largest in winter and early spring. Boxplots show monthly size distributions; sample sizes (n) are indicated above each box. White diamonds represent means, and points are jittered for visibility. (b) Monthly mean radius (± 95 % CI) of all sea stars within a frame (black circles, solid line) compared with asymptomatic individuals only (orange triangles, dashed line). Both groups showed a decline in mean size during summer followed by partial recovery in autumn. (c) Seasonal trend in sea star density (individuals m^2^), estimated using a negative-binomial GLMM with a temporal spline (± 95 % CI), showing a pronounced summer peak followed by an autumn decline.

**Figure 4.**
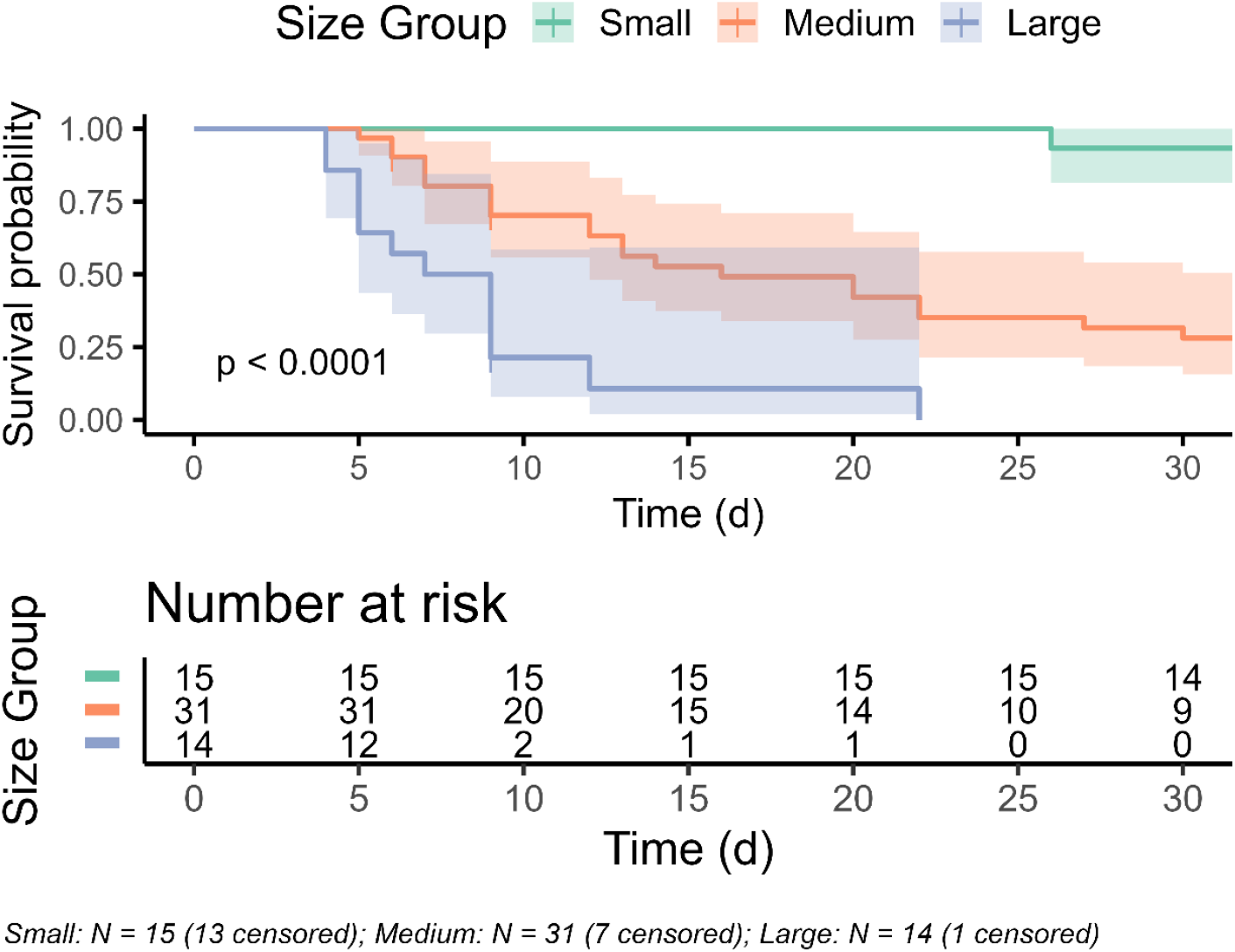
Kaplan-Meier survival curves for sea stars grouped by body size over a 33-day period. Larger individuals showed lower survival than smaller ones. Shaded areas represent 95% confidence intervals. Group sizes: Small (< 4.0 cm; n = 15, 13 censored), Medium (4.0 - 6.5 cm; n = 31, 7 censored), Large (> 6.5 cm; n = 14, 1 censored). Survival differences were significant (log-rank test, *p* < 0.0001).

Sea star density varied strongly over the year, with a clear summer peak (29.3 ind. m^-2^ in July) followed by a decline into autumn. Terminal disease cases showed similarly dynamic patterns. Interval-based mortality estimates ranged from ~70 to >500 individuals per week within the surveyed reef area (Supplementary Fig. S1), indicating temporal variability in disease associated mortality.

### 3.4 Size-dependent survival under laboratory conditions

Body size differed significantly among treatment groups, with larger individuals more frequently found in the Diseased group (arm length: ANOVA, F_2_,_57_ = 14.79, *p* < 0.001; wet weight: F_2,57_ = 7.05, *p* = 0.0018). This imbalance reflected the scarceness of large, asymptomatic sea stars in the field, and the lack of effect of the exposure treatment (only one individual developed symptoms). Disease progression varied by treatment (Kruskal-Wallis χ^2^= 17.92, *p* < 0.001), but a Cox proportional-hazards model showed that these differences were fully explained by body size. After adjusting for arm length and weight, treatment no longer significantly affected survival (*p* > 0.1). In contrast, arm length remained a strong predictor of mortality risk (HR = 2.47, *p* < 0.001), whereas wet weight had no significant effect (*p* = 0.32). Categorical size models confirmed higher mortality in medium- (HR = 12.6, *p* < 0.001) and large-sized individuals (HR = 50.8, *p* < 0.001) compared with small ones, and a log-rank test supported significant survival differences among all size classes (χ^2^= 42.8, *p* < 0.0001). The final model showed strong predictive performance (concordance = 0.83), indicating that body size, rather than treatment, was the main determinant of survival.

## 4 Discussion

### Field monitoring reveals a persistent wasting disease in Baltic *Asterias rubens*

In this study, we used year-round field monitoring to characterise the occurrence, severity, and temporal dynamics of sea star wasting symptoms in *A. rubens* from the western Baltic Sea. Our results reveal a persistent wasting-type disease is affecting a substantial fraction of the population across seasons. By combining field prevalence data with laboratory assays, we further show that disease expression and mortality are strongly size-dependent, disproportionately affecting large adult individuals. Together, these findings indicate that wasting in Baltic *A. rubens* represents an established disease with the potential to alter population structure and ecological function.

### Seasonal prevalence and symptom expression in Baltic *Asterias rubens*

Across all monitored sites and seasons, a large fraction of the monitored population consistently exhibited wasting symptoms consistent with SSDW, including skin softening, arm curling, pale lesions, open wounds, body-wall disintegration and autotomy [2,5,10,14]. Prevalence remained consistently high throughout the year, with seasonal variability showing comparatively higher values in early spring and early autumn and lower but still substantial levels in winter and mid-summer. Even at its annual minimum, prevalence rarely dropped below ~35-40%, indicating that wasting symptoms are persistently established in the population rather than short-lived episodic events. While these values do not match the catastrophic mass mortality seen in the Pacific during 2013-2015, they indicate that a wasting-type disease is prevalent in the Baltic at a scale that warrants attention.

Sea star wasting disease has a long and well-documented history in the northeastern Pacific, with descriptions extending from the late 19th century through repeated events in the 1970s-1990s to the major 2013-2015 outbreak [2,3,6,8,14,28,61–63]. At peak disease periods, prevalence in keystone species such as *Pisaster ochraceus* and *Pycnopodia helianthoides* frequently reached 60-100%, and *P. helianthoides* populations declined by approximately 90%, with surviving individuals largely restricted to juveniles [28,61]. Mild lesions dominated early outbreak stages [27], and long-term data now show that lesions remain the most common symptom years after the initial collapse [64].

In contrast, field observations of wasting-like symptoms in Europe have been sparse, limited to reports of mortality and lesions in *Astropecten jonstoni* along the Sicilian coast between 1999 and 2003 [5] and reports from aquaria and research facilities describing arm curling, bulging, lesion formation and arm loss in captive *Asterias rubens* and *Crossaster papposus*, but without environmental context [11–13]. Although SSWD-like symptoms have been reported, the absence of systematic field monitoring has so far prevented reliable assessment of disease prevalence, persistence, and population-level impacts in European sea star populations.

Some historical and experimental evidence suggests that wasting-like symptoms may have occurred in Baltic *A. rubens* well before the present study. A 2016 laboratory experiment on individuals from Sylt and Kiel Fjord documented rapid mortality characterised by white lesions, arm autotomy, and tissue degradation in both control and treatment groups, closely resembling SSWD [65]. Scientific divers have also reported occasional observations of degraded sea stars during warm summers (F. Melzner, Martin Wahl pers. comm.), suggesting that such events were sporadic but largely overlooked. Early-stage SSWD in *A. rubens* can resemble osmotic or thermal stress responses, and echinoderms are known to autotomise under environmental stress [66]. The limited reporting of wasting-like symptoms in Europe therefore likely reflects limited monitoring effort rather than true absence. Together, these observations raise the question of whether wasting-like patterns observed across different regions reflect a single disease or a broader syndrome complex with multiple possible causes.

### Wasting disproportionately affects larger, adult *A. rubens* in the Baltic

The summer decline in SSWD prevalence coincided with a reduction in the average size of monitored sea stars. Given the strong size-dependent mortality observed in both field surveys and laboratory assays, this midsummer minimum most likely reflects the selective loss of large, highly susceptible individuals, leaving behind smaller juveniles and recent recruits. Over the same period, sea star density per m^-2^ increased from spring into early summer before declining again toward autumn. This increase in density, despite ongoing adult mortality, indicates that strong recruitment temporarily masked the loss of larger individuals.

Similar patterns have been reported in other asteroid species affected by SSWD, including the post-outbreak surge of juvenile *Pisaster ochraceus* following mass mortality events in 2014-2016 [8] and recruitment pulses documented in several additional studies [5,7,64]. Within the analysed subset of the monitored reef area, 3,250 individuals were recorded in terminal disease categories. Scaled to the full reef area, this corresponds to approximately 30,000 terminal cases (~16% of the population) observed during the sampling period. This estimate represents a minimum, as it is based on seven discrete surveys and reflects only the standing stock of moribund individuals at the time of observation. If terminal individuals typically die within one to two weeks, a substantial proportion of mortality likely occurred between surveys and remained undetected. Therefore, cumulative annual mortality may exceed our point estimates by several-fold. Because disease progression and mortality vary seasonally and among individuals, we do not apply a fixed correction factor and instead emphasise the uncertainty surrounding absolute mortality rates. Nevertheless, these estimates indicate substantial SSWD-associated mortality that disproportionately removed larger individuals.

Seasonal variation in SSWD prevalence further supports a mechanistic role of environmental stressors that disproportionately affects larger sea stars. In winter, prevalence was lowest (~35% compared with ~55% in spring and autumn), consistent with evidence from other regions that cold, saline conditions can reduce wasting severity [5,28,31,67]. In contrast, spring and summer conditions impose multiple interacting stressors, including episodic salinity fluctuations, stratification-driven hypoxia, and frequent upwelling events [39,40,68] that are known to exacerbate physiological strain in *A. rubens* [49,52,69]. These patterns align with known physiological constraints in *A. rubens*: Larger sea stars have a lower surface-area-to-volume ratio, reducing diffusive oxygen uptake while simultaneously having higher metabolic demand [16,70,71]. Under thermal, osmotic, or metabolic stress, this combination limits resilience and increases susceptibility to tissue damage and infection - a pattern also reported in other invertebrates such as oysters and abalones [24,25,72,73]. Further, seasonal reproduction may amplify this size-selective vulnerability. In the Baltic Sea, *A. rubens* replenishes its gonads through winter and spawns in spring, an energetically demanding process largely restricted to individuals above ~5 cm radius [74,75]. Although spawning itself was not a significant predictor of mortality in our laboratory assay (likely because animals were held under stable conditions), post-spawning energetic depletion may increasing mortality risk for large adults [24,26]. Together, size-selective mortality and seasonal stress exposure indicate that wasting does not affect all individuals equally but mostly affects large adults, with consequences for population size structure and recovery potential.

### Ecological implications

As the dominant echinoderm predator in the western Baltic, *A. rubens* exerts strong top-down control on blue mussels (*Mytilus* spp.) [45]. Baltic populations already experience chronic osmotic stress due to frequent salinity fluctuations between ~11 and 20, which contribute to their smaller body size, reduced calcification, and shorter spawning period compared with North Sea populations [76]. The decline in mean body size and the elevated mortality of large individuals observed here may therefore alter predation pressure on mussel beds, as preferred mussel size scales with sea star size [77,78]. Shifts in predator size structure can modify benthic communities, as shown in Pacific systems where SSWD-driven predator loss caused an 80% decline in kelp coverage due to increased prey abundances [61]. In the Baltic, a comparable risk exists for eelgrass meadows. Blue mussel expansion has been shown to suppress eelgrass through overgrowth and competition, meaning that reduced sea star predation could indirectly contribute to the loss of eelgrass meadows as a key habitat-forming ecosystem supporting high biodiversity [46].

Recovery from SSWD-associated losses may be further constrained by the already unstable environmental conditions of the Baltic Sea. Recruitment success and early growth of *Asterias rubens* are highly sensitive to salinity, with larval development failing at ≤12 and settlement in Kiel Fjord occurring only during years of high and stable salinity [49,74]. In addition, both summer heatwaves and winter warming impose energetic constraints by increasing metabolic demand while reducing feeding activity and energy availability, thereby slowing growth and delaying recovery in both juveniles and adults [48,50,52]. When combined with projected long-term warming and desalination trends in the western Baltic [37], these factors are likely to narrow the temporal window for successful recruitment and growth. Additionally, echinoderm populations are known to exhibit strong fluctuations in abundance and size structure, driven by episodic recruitment pulses and mortality events (reviewed by [79,80]. The ecological significance of such changes is often difficult to assess because long-term baseline data needed to define a “natural” population state are rarely available. This limitation is particularly relevant for *A. rubens* in the Baltic Sea, where early studies already reported pronounced variability in abundance and size-class structure driven by hydrodynamics, variable food availability, and temperature mediated differences in growth, complicating interpretation based on density or size alone [74,81]. However, continuous monitoring over longer timescales is largely missing for this system.

In this context, the high recruitment observed following periods of elevated mortality in our surveys should not be interpreted as population recovery. Recruitment-driven increases in *Asterias* populations are often episodic and transient [79]. Consistent with this, long-term studies from the northeastern Pacific show that following SSWD, population density and biomass can recover through recruitment, while average body size remains substantially reduced for many years due to the persistent loss of large adults [64]. Our findings suggest that a similar pattern may be emerging in Baltic *A. rubens*, where repeated, size-selective mortality could maintain a persistently younger and smaller population structure despite high recruitment. Larger individuals typically contribute disproportionally to reproductive output, meaning that the loss of large adults may reduce the population reproductive capacity even in years with strong recruitment, a pattern well known from exploited fish populations such as Baltic Cod [82,83]. Continued long-term monitoring will therefore be essential to distinguish natural population variability from sustained, disease-driven change and to assess the long-term consequences for population resilience and benthic community dynamics.

### Is there only one Sea Star Wasting Disease?

Wasting-like symptoms including lesions, arm curling, softening, bulging, and rapid tissue disintegration, have been reported from more than 20 asteroid species worldwide, across both natural populations and captive settings [1,10,12–14,84,85]. While a specific bacterial agent has recently been identified as causative for wasting in *Pycnopodia helianthoides* [22], the broad taxonomic and geographic spread of similar symptom trajectories makes it unlikely that all wasting events arise from a single pathogen or uniform mechanism [32,64]. Instead, accumulating evidence suggests that multiple microbial agents, together with environmentally mediated physiological stress, can produce comparable disease phenotypes across species and regions.

This interpretation is supported by repeated associations between wasting severity and elevated temperature, reduced oxygen availability, organic matter enrichment, and shifts in the sea star-associated microbiome [17,19–21,84,86]. Several studies have documented microbial dysbiosis in diseased individuals, and in some cases even in apparently healthy sea stars prior to overt symptom development, suggesting that wasting may reflect a breakdown of host-microbe equilibrium rather than infection by a single obligate pathogen [20,21,86]. Within this framework, different stressors may affect similar downstream processes such as impaired tissue oxygenation [17], immune dysfunction [21], or microbial overgrowth and thereby ultimately producing the characteristic wasting phenotype. In our experiment, wasting symptoms in *A. rubens* occurred even under experimentally stable conditions, indicating that susceptibility and outcome are shaped by host condition and prior exposure rather than by acute environmental extremes alone. Together, these findings support the view that SSWD represents a complex, context-dependent disease syndrome, the expression of which may vary across species, environments, and physiological states. While a specific causative agent cannot be excluded in Baltic *A. rubens*, the relative roles of microbial pathogens, host condition, and environmental stressors remain unresolved. Disentangling these mechanisms will require targeted microbial screening and experimental approaches, which are currently lacking for Baltic populations.

## Conclusion

Across all surveys, approximately 20% of *Asterias rubens* exhibited severe wasting symptoms, and an additional ~25% showed consistent early-stage signs such as arm curling and bulging. Together with the laboratory assay, our field observations show that the disease disproportionately affects larger individuals, leading to repeated loss of adults and pushing the population toward a persistently smaller size structure despite strong recruitment. This study provides the first continuous, field-based assessment of wasting disease in European *A. rubens* and demonstrates a shift from historically sporadic observations to widespread, sustained prevalence and mortality. Identifying the microbial agents involved and experimentally resolving if temperature, salinity fluctuations, hypoxia and organic matter loading interact with disease processes remain important next steps. Long-term monitoring across European sites will be essential for determining whether recurring wasting, recruitment constraints and ongoing environmental change will alter the ecological role of *A. rubens* strongly enough to reshape mussel bed dynamics and benthic community structure. Our findings highlight a disease dynamic in the western Baltic that warrants careful attention, both for the future of this keystone predator and for the stability of the ecosystems it inhabits.

## Acknowledgements

We thank the interns Rieke Lühring, Emily Jacobsen, Svea Jonas, Janne Held, Shelby Lynn and Theresa Lütt for their assistance with data collection, even under challenging weather conditions. We gratefully acknowledge the support of technicians Björn Buchholz and Fabian Wendt for their help with the camera setup. We also thank the CAU diving team for their efforts in assisting with the collection of sea stars.

## Author Contributions

Katja Seebass: Conceptualization, methodology, investigation, formal analysis, writing-original draft, review and editing. Frank Melzner: Conceptualization, methodology, writing - review and editing. Vinzent Ferfers: Investigation (fieldwork and data collection), data curation. Jahangir Vajedsamiei: Formal analysis (statistical analyses), visualisation. All authors approved the final version of the manuscript.

## Data Availability

Data supporting the findings of this study are not yet publicly available, as analyses are ongoing, but will be made available upon reasonable request and in a future peer-reviewed version of this manuscript.

